# FlowBench: separating planning, fault recovery and interpretation in agentic bioinformatics

**DOI:** 10.64898/2026.06.12.731844

**Authors:** Alina Kurjan, Adam P. Cribbs

## Abstract

Agentic large language model (LLM) systems are being deployed in bioinformatics faster than they are understood, and single-metric evaluations conflate capabilities that fail independently. We introduce FlowBench, a benchmark that decomposes agentic bioinformatics performance into planning, fault recovery, biological interpretation, and end-to-end output-fidelity. Existing systems achieve high plan completeness, but their closed, single-provider designs prevent attribution of performance to scaffolding versus the underlying model. We therefore built FlowAgent, a modular, provider-agnostic framework whose components can be selectively disabled and whose backbone model can be swapped across providers on a shared harness, and used it to evaluate 23 models from three main providers. Three findings emerge. First, generating a valid workflow plan from a named toolchain is largely solved, whereas inferring an appropriate toolchain from biological intent alone is uniformly difficult regardless of model tier, compressing all models into a narrow 44–57% pass-rate band. Second, ablation shows that the dependency-structured plan and a completeness-reflection step drive performance, while adding a same-context validator-driven retry makes structural quality worse. Third, fault recovery and data-grounded interpretation remain unsolved. Models frequently propose fixes that force a clean exit while leaving the underlying data invalid, and data-grounded interpretation lags internal-knowledge recall by a consistent margin. Safety does not emerge from capability, and reasoning-tier models were among the least reliable at recognising unrecoverable faults. Once planning saturates, agent architecture and refusal calibration, not model scale, are the productive frontier.

**Availability and implementation:** FlowAgent and FlowBench are available under a GPLv3 licence at https://github.com/EnteloBio/flowagent

## 1 Introduction

Agentic large language model (LLM) systems that translate natural-language requests into executable bioinformatics pipelines are rapidly moving from proof-of-concept to deployment. Systems such as Biomni huang2025biomni, AutoBA zhou2023autoba, ChemCrow bran2024chemcrow, BioMaster su2025biomaster, BioAgents mehandru2025bioagents and CellAgent xiao2024cellagent allow scientists to describe an analysis in plain English and receive a structured pipeline in return, promising to lower the programming barriers that have long limited access to complex workflows spanning single-cell RNA sequencing, spatial transcriptomics, methylation, chromatin conformation, and metagenomic analyses shade2015. Traditional workflow management systems such as Nextflow ditommaso2017, Snakemake koster2012, CGAT-core cribbs2019, and Galaxy galaxy2024 address reproducibility and portability but demand programming expertise, which further motivates natural-language interfaces. Because each system has been evaluated in isolation, with different metrics, prompt sets and model backends, the field lacks answers to basic questions about when and why these systems succeed or fail.

These systems are each presented behind a closed interface that couples one architecture to a single model family, so a ranking of finished products cannot explain why one outperforms another. The contribution of the scaffolding cannot be separated from that of the model, a component cannot be removed to measure its effect, nor the architecture held fixed while the model is changed. Answering such questions requires a system that can be taken apart and run on different models under identical conditions. We built FlowAgent for this purpose philpott2025flowagent, a modular, provider-agnostic framework in which the backbone model and each evaluated planner component can be toggled independently and run across OpenAI, Anthropic and Google Gemini models on a shared harness. Beyond making architecture and model separately testable, an open framework also exposes the cluster and HPC backends that a closed local tool does not.

FlowAgent’s planner emits, by default, a dependency-structured workflow plan (Supplementary Figure 1), with nodes representing analytical tools and edges encoding input–output dependencies, and the resulting step graph is checked for acyclicity. In formal terms this is a domain-specific causal representation. FlowAgent makes explicit the ordering constraints that an unstructured agent would otherwise hold implicitly, reflecting a broader convergence between structured knowledge representations and LLMs across AI research pan2023, bran2024chemcrow, mavromatis2024, bian2025. Whether this structure does real work, rather than the underlying model, is an empirical question, and a controllable framework can answer it through ablation.

Here we present FlowBench (Figure 1), a reproducible benchmark that decomposes agentic bioinformatics into four layers: planning, fault recovery, biological interpretation, and a planned end-to-end output-fidelity evaluation. Planning is assessed on 66 prompts across three regimes. Recovery is assessed on a 28-fault catalogue across three difficulty tiers, and interpretation on questions anchored to public datasets and stratified by the kind of evidence each requires. Three findings emerge. First, generating a structurally valid plan from a named toolchain is largely solved, whereas selecting a toolchain from biological intent alone is not. Second, controlled ablation shows that enforced dependency structure and a completeness-reflection step, not model scale, drive plan quality. Third, fault recovery and data-grounded interpretation remain the genuine open problems.

**Figure 1.**
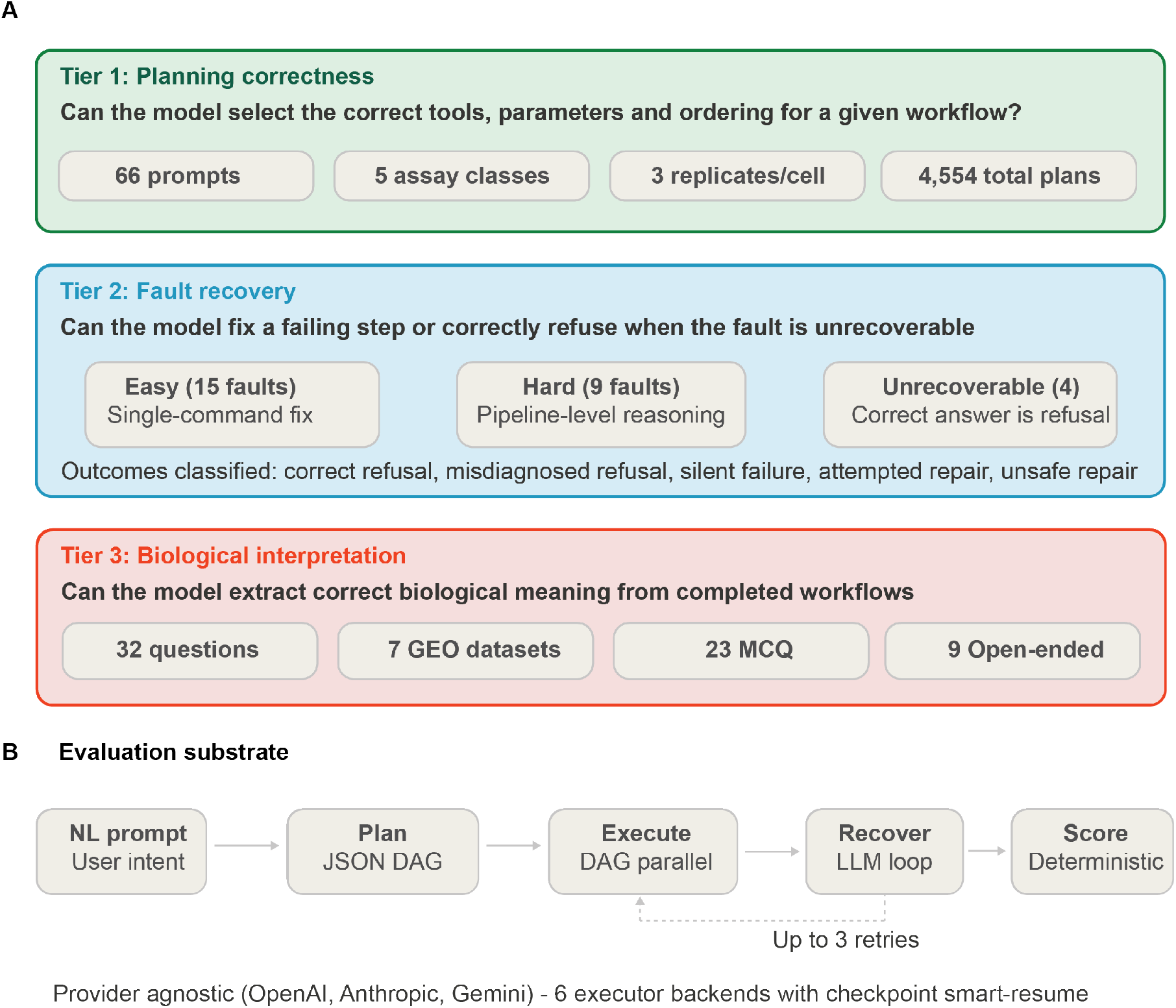
FlowBench three-tier evaluation framework and FlowAgent execution substrate. Overview of the benchmark stack (top) and runtime path (bottom). Tier 1 (green), planning correctness: models receive natural-language bioinformatics requests and emit a structured workflow plan; the corpus comprises 66 prompts across five assay classes (23 standard explicit-tool, 18 hard explicit-tool, 25 tool-inference) at three replicates per model–prompt cell (4,554 plans). Tier 2 (blue), fault recovery: 28 injected faults across three severities (15 easy single-command fixes, 9 hard pipeline-level faults, 4 unrecoverable), with outcomes classified as correct refusal, misdiagnosed refusal, attempted repair, silent failure, or unsafe repair. Tier 3 (red), biological interpretation: 32 questions over 7 GEO-derived datasets (23 multiple-choice, 9 open-ended). Bottom: the FlowAgent runtime maps user intent to a JSON DAG plan, executes the DAG in parallel, enters an LLM-driven recovery loop (up to three retries per failed step), and applies deterministic scoring. Backends are provider-agnostic (OpenAI, Anthropic, Google) across executors with checkpoint smart-resume.

## 2 Results

### 2.1 Closed agents plan well but cannot be ablated, motivating an open evaluation framework

We first benchmarked the existing agentic systems on two complementary measures of planning quality: plan correctness, the fraction of prompts whose generated plan passes the full rubric (Figure 2A), and tool-completeness (i.e. the fraction of rubric-required tools each system places correctly) (Figure 2B). Among the published bioinformatics agents, completeness was uniformly high, with BioMaster, AutoBA and Biomni reaching 87%, 83% and 70%, respectively, but plan correctness was the more discriminating measure. The same systems passed roughly 99%, 90% and 82% of prompts. Claude Code, a general-purpose agentic coding tool with no bioinformatics-specific design, outperformed all three dedicated agents on both measures, reaching 100% completeness and near-perfect plan correctness. Failures in the lagging systems were predominantly planning errors rather than execution faults. Plan-level failures accounted for the shortfall in every case (3, 4 and 9 calls for AutoBA, BioMaster and Biomni), and only AutoBA produced upstream execution crashes (Figure 2C).

**Figure 2.**
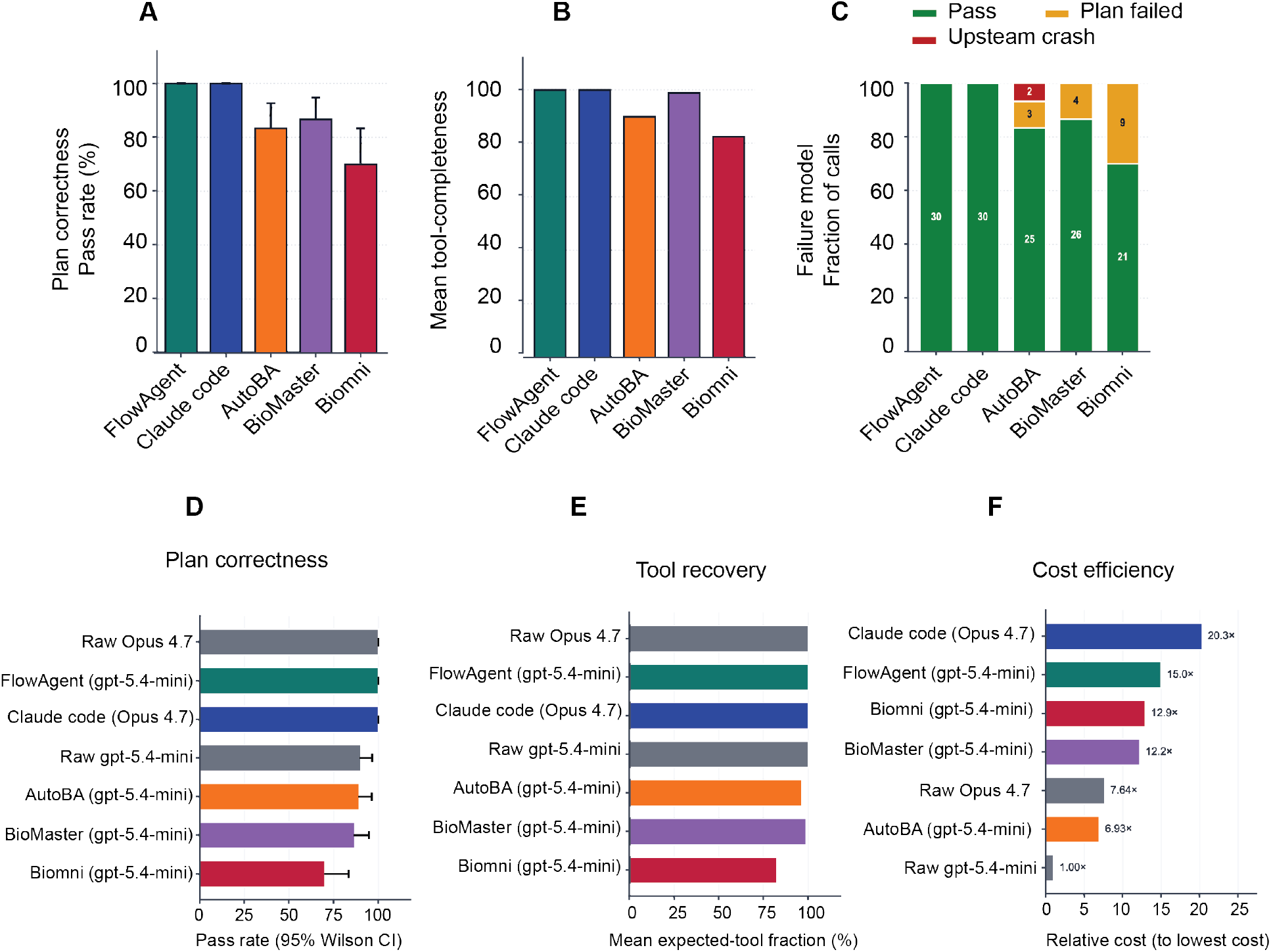
Planning performance, accuracy, and cost across prompt tiers for a representative five-model subset. Five models evaluated on the FlowAgent planning benchmark under three prompt regimes. (A) Pass rate on standard explicit-tool prompts (tools named in the request). (B) Pass rate on hard explicit-tool prompts (niche assays, tools still named). (C) Outcome composition on tool-inference prompts (goal and input data only, no tool names): correct plan (green), incorrect tool choice (yellow), incorrect workflow type (red); bar height is 30 prompts per model, with counts annotated. (D) Aggregate pass rate across the full corpus. (E) Planning accuracy (structural and rubric adherence). (F) Relative cost per successful plan, normalised to the cheapest model (1.00 *×*). Bars in A, B and E carry replicate-derived error bars; segments in C are stacked counts.

Because Claude Code is a closed, single-vendor product, its scaffold and backbone model cannot be separated, and benchmark rankings alone cannot establish whether performance derives from one, the other, or both. To address this need, we developed FlowAgent, an open, model-agnostic framework whose components can be selectively disabled and whose backbone model can be swapped across providers on a shared harness philpott2025flowagent. As a baseline, FlowAgent matched Claude Code on both measures (100% completeness and near-perfect plan correctness; Figure 2A–C).

To separate the contribution of the backbone model from that of the scaffolding, we re-ran all systems wrapped with a common `gpt-5.4-mini` backbone and compared them against their raw, unwrapped models as references (Figure 2D–F). Keeping the backbone constant removed most of the apparent advantage of the dedicated bioinformatics agents. AutoBA and BioMaster reached approximately 88% and 87% plan correctness, respectively, with neither exceeding raw gpt-5.4-mini (approximately 88%), while Biomni fell below it (approximately 70%), indicating that dedicated-agent scaffolding adds little over the unwrapped model and in one case even degrades it. FlowAgent was the exception, raising the same backbone to 100% plan correctness, equivalent to raw Opus 4.7. Reading the ranking from the top, raw Opus 4.7 alone matched Claude Code’s near-perfect correctness, suggesting that Claude Code’s planning may be attributable to its underlying Opus model rather than its coding scaffold. Tool recovery was uniformly high across both raw and wrapped systems, with only Biomni lagging, confirming that plan correctness rather than tool retrieval is the discriminating axis. These results came at sharply different costs (Figure 2F): raw `gpt-5.4-mini` was cheapest by construction and Claude Code the most expensive, while FlowAgent reached near-Opus correctness on the cheaper backbone at intermediate cost, below Claude Code but above the raw Opus baseline.

### 2.2 Executed pipelines reproduce published RNA-seq differential expression, dissociating plan correctness from scientific reproduction

Establishing FlowAgent as a measurement tool requires more than parity on planning; its executed pipelines must reproduce real biology, not merely produce plausible plans. We therefore compared FlowAgent-derived differential expression (DE) tables (generated by genuine download, quantification and DE analysis rather than mock execution) against curator-built reference tables from published RNA-seq studies (Figure 3). Gene-level concordance was high across heterogeneous GEO cases (Spearman *ρ* up to 0.98 on log_2_ fold change; Jaccard overlap of the top-200 reference DEGs up to 0.59), confirming that the harness executes faithful end-to-end workflows and that its plan-level measures are anchored to reproducible scientific output. The stratification further shows that agreement with published numbers depends on execution-time tool choice, rather than rubric compliance alone, dissociating plan correctness from scientific reproduction and confirming that FlowAgent can run, not only plan.

**Figure 3.**
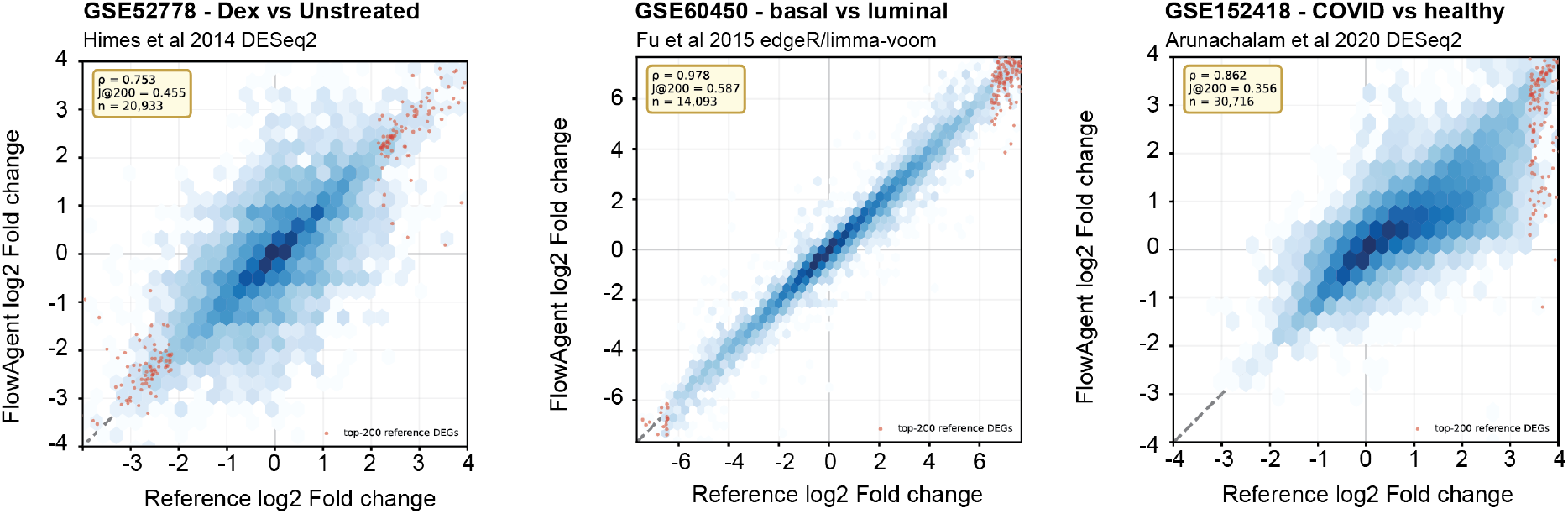
Output fidelity of agent-generated differential expression, stratified by planning prompt tier. Hexagonal density plots of reference log_2_ fold-change (*x*-axis) against agent log_2_ fold-change (*y*-axis) for genes passing expression filters; darker bins denote higher gene density. The dashed diagonal marks perfect agreement (*y* = *x*), and red points highlight the top-200 reference differentially expressed genes (DEGs). Inset statistics report Spearman *ρ*, the Jaccard overlap of the top-200 DEG sets (J@200), and the number of genes plotted (*n*).

#### Planning correctness, cost and task difficulty across a bioinformatics workflow benchmark

Using FlowAgent as the evaluation scaffold, we tested 23 models from three providers (Anthropic, OpenAI and Google) on an automated planning-correctness benchmark built from 66 realistic bioinformatics prompts, each run in triplicate, spanning three difficulty regimes (Figure 4, Supplementary Figures 2–5). Standard explicit-tool prompts (*n* = 23) cover everyday workflows by naming the expected software (e.g. kallisto, STAR or MACS2), and are scored on the workflow type, full coverage of the named tools, exclusion of forbidden tools, and a minimum step count. Hard explicit-tool prompts (*n* = 18) apply the same rubric to niche or multi-step assays (such as bisulfite sequencing, metagenomics, Hi-C, long-read assembly and differential binding) where tool names are provided; however, the difficulty lies instead in longer chains, domain-specific choices and R-package wrappers. Finally, tool-inference prompts (*n* = 25) give only the biological goal and input data, never naming a tool, so the model is expected to select an appropriate pipeline rather than transcribe supplied names. Each is paired with a pre-registered set of acceptable, functionally equivalent toolchains (for example STAR with featureCounts, or salmon with tximport), and a plan passes only if it reproduces at least one set in full, with each required tool invoked as the executable at the head of a well-formed command, crediting genuine toolchain selection rather than tool names listed in valid JSON.

**Figure 4.**
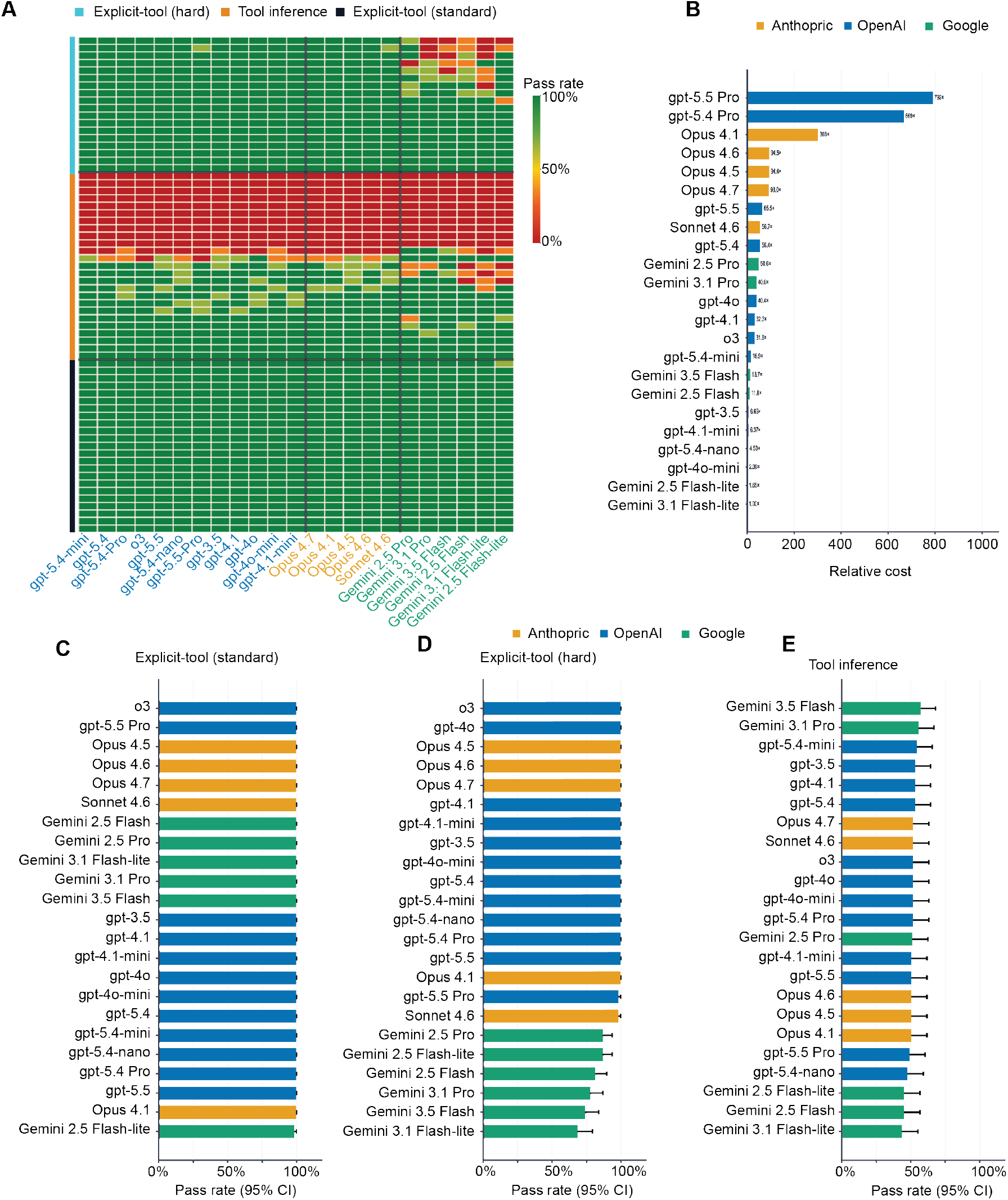
Planning correctness and cost for 23 LLMs on the 66-prompt FlowAgent corpus. Models from Anthropic, OpenAI and Google, each producing a JSON workflow plan per prompt over three replicates (aggregate pass rates shown). Scoring rubric: workflow type, tool coverage (explicit-tool tiers) or realisation of an acceptable toolchain (inference tier), forbidden-tool exclusions, and minimum step count. (A) Per-prompt by per-model heatmap of pass rate (green, pass; red, fail). Rows are grouped by corpus tier (standard explicit-tool, tool-inference, hard explicit-tool), indicated by the left-hand row colour; columns are coloured by provider (Anthropic, orange; OpenAI, blue; Google, green). The contiguous red band corresponds to the tool-inference tier and persists across all models. (B) Mean cost per prompt (USD), expressed relative to the cheapest model (1.00*×* to 792*×*). (C) Pass rate on standard explicit-tool prompts (*n* = 23). (D) Pass rate on hard explicit-tool prompts (*n* = 18). (E) Pass rate on tool-inference prompts (*n* = 25); error bars are 95% Wilson confidence intervals. Model ordering is consistent across panels within each provider family.

The results reveal a sharp split by task type (Figure 4A). On standard explicit-tool prompts, pass rates were uniformly high and most models scored at or near 100% (Figure 4A,C). On hard explicit-tool prompts, frontier OpenAI and Anthropic models remained near-perfect, while several Google models fell slightly below (Figure 4A,D), with failures confined to a few niche assays such as differential-binding and long-read assembly. Tool-inference prompts told a different story: all models compressed into a narrow *∼*44–57% band with overlapping confidence intervals (Figure 4A,E). A contiguous block of these prompts failed across all models due to incorrect toolchain selection rather than formatting errors. No model approached its explicit-tool performance on this regime, confirming that selecting an appropriate toolchain from biological intent alone, rather than plan generation under explicit instructions, is the dominant bottleneck.

Planning cost spanned roughly three orders of magnitude relative to the cheapest model (Gemini 3.1 Flash-Lite; Figure 4B and Supplementary Figure 6). Lightweight Google models sat at the low end (*∼*1–2*×*) and mid-tier OpenAI and Anthropic models in the middle (*∼* 5–65*×*), while flagship and reasoning-tier models were far more expensive (Opus 4.1 *∼* 303*×* , GPT-5.4 Pro *∼*669 *×*, GPT-5.5 Pro *∼*792*×*). Since explicit-tool planning performance was already near ceiling for most models, and tool-inference performance was similarly plateaued across all of them, cost was not correlated with accuracy on either task component.

### 2.3 Data-grounded interpretation is the consistent weak point across all models

We assessed 11 Anthropic, OpenAI and Google models on a multiple-choice question (MCQ) and open-ended question task probing biological interpretation (chance = 25%). Overall MCQ accuracy (Figure 5A), using gpt-5.4 as the judge model (Supplementary Figure 7), compressed the models into a narrow *∼*75–80% band, with `gpt-5.4` and`gpt-5.5-pro` highest (80%) and a tightly overlapping cluster of Claude and Gemini models at 72.5–77.5%, with confidence intervals overlapping broadly throughout. Per-dataset accuracy (Figure 5B) was high and consistent across most cases (78–100%), with two exceptions: the COVID blood differential expression dataset (`gse152418_covid_blood_de` , 40%) and the mammary dataset (`gse60450_mammary_de` , 50%) were uniformly difficult across all models, localising the performance deficit to specific biological contexts rather than distributing it evenly across the benchmark. Open-ended answer quality (Figure 5C and Supplementary Figure 8) mirrored this pattern, with mean judge scores clustered tightly at 73–77/100 with overlapping standard deviations, and no model separated clearly from the field.

**Figure 5.**
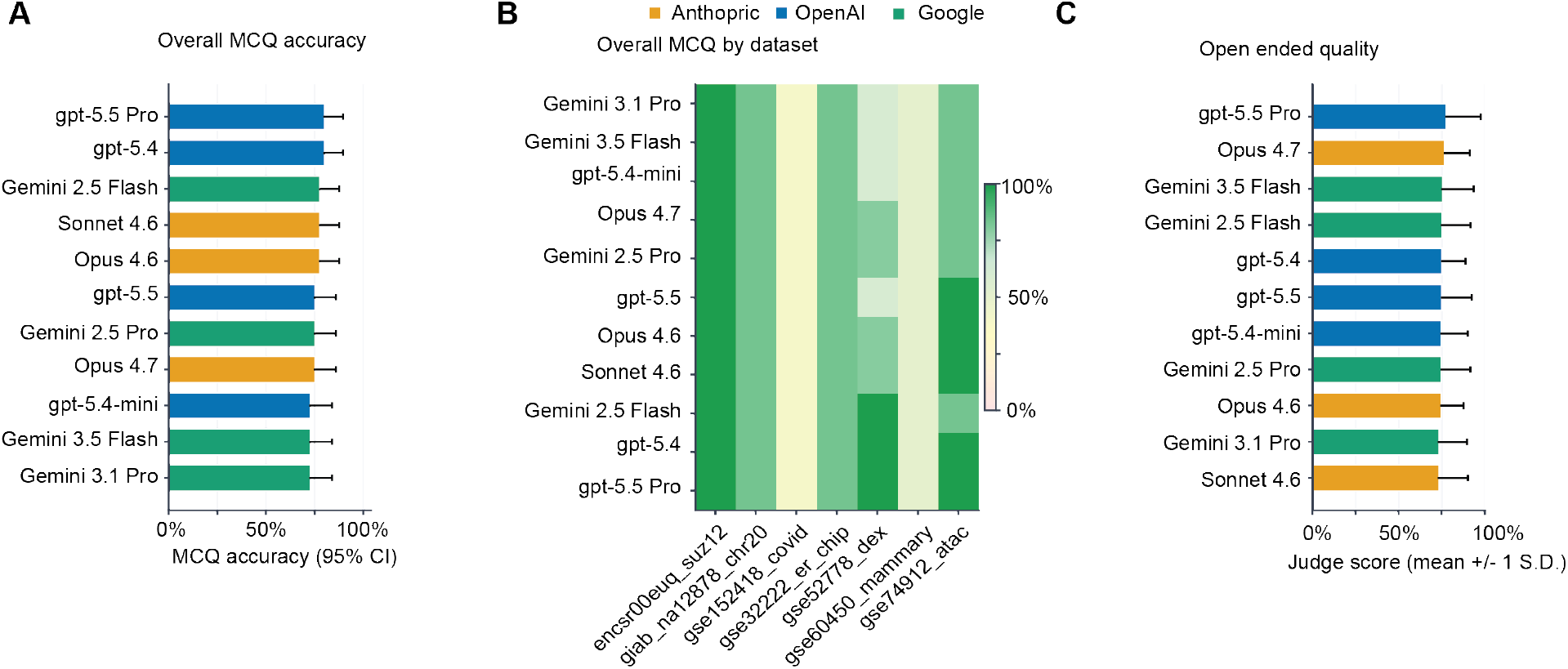
Biological-interpretation performance across 11 models. Eleven models from Anthropic, OpenAI and Google evaluated on the FlowBench interpretation benchmark, which supplies the plain-text outputs of completed workflows and poses multiple-choice and open-ended questions (chance = 25%), scored by an LLM judge (`gpt-5.4`). Bars are coloured by provider (Anthropic, OpenAI and Google). (A) Overall MCQ accuracy per model; values compress into a narrow band of approximately 75 to 80%, with `gpt-5.4` and `gpt-5.5` Pro highest (80%) and the Claude and Gemini models clustered at 72.5 to 77.5%. (B) Per-dataset MCQ accuracy as a model-by-dataset heatmap over the seven public datasets, with colour encoding accuracy (darker green higher, pale tones the lowest-scoring datasets); agreement is high and consistent across most datasets (78 to 100%) and falls in two uniformly difficult cases: the COVID blood differential-expression dataset (`gse152418_covid_blood_de` , 40%) and the mammary dataset (`gse60450_mammary_de` , 50%). (C) Open-ended answer quality, the mean LLM-judge score (0 to 100) per model with error bars; scores cluster tightly at 73 to 77 with overlapping spread, and no model separates from the field.

### 2.4 Dependency structure and structured reflection, not model scale, drive plan quality

To determine which architectural components drive planning performance, we built FlowAgent as a modular framework whose components can be selectively disabled. We ablated five components of the FlowAgent planning stack in paired ON/OFF comparisons (Figure 6; Table 1; Supplementary Figure 9), scoring each plan on two independent axes. *Overall pass* measures correctness against the benchmark rubric (valid schema and directed acyclic graph (DAG), correct workflow type, all expected tools present, no forbidden tools, and sufficient steps), whereas *completeness pass* measures the internal structural quality of the workflow graph (alignment steps with an index or download ancestor, no dangling downloads, quantification and analysis steps reaching a terminal sink, and a weakly connected graph). Completeness is recorded alongside overall pass but does not gate it, so the two can diverge: a plan can pass the rubric with a broken graph, or fail it while remaining structurally sound.

**Figure 6.**
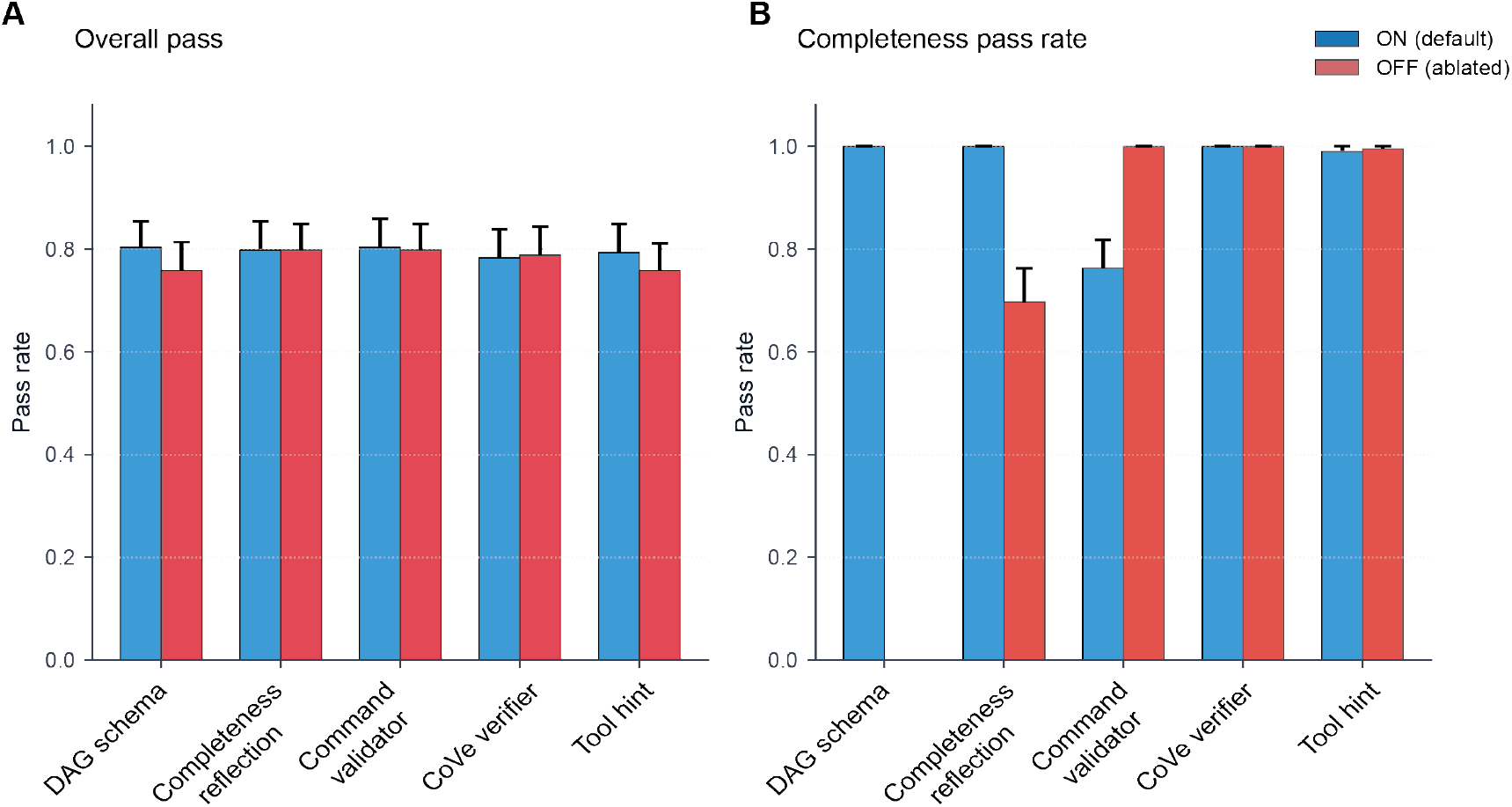
FlowAgent planning-stack ablation. Each of five components was toggled ON (blue) versus OFF (red) while the others were held at defaults (`gpt-5.4-mini` , 66 prompts, three replicates). (A) Overall pass rate. (B) Completeness pass rate. Error bars are 95% Wilson confidence intervals; McNemar *p*-values are reported in Table 1.

**Table 1:**
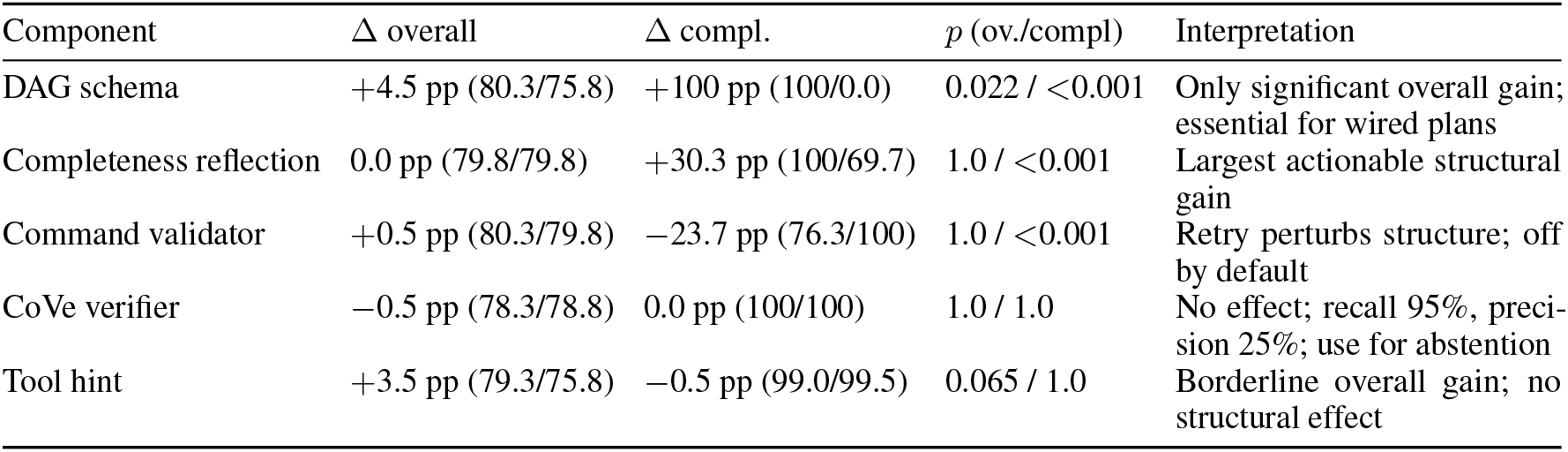
Paired ON/OFF ablation of five FlowAgent planning-stack components. Each component was toggled on and off while the others were held at their defaults, and the same prompts were scored under both settings (`gpt-5.4-mini` , 66 prompts *×* 3 replicates, *n* = 198 paired plans per component). Overall pass measures correctness against the benchmark rubric; completeness pass measures the structural quality of the workflow graph. Δ columns give the change on enabling each component (pp = percentage points; positive = improvement). *P* -values from McNemar’s exact test on discordant pairs are reported for overall pass and completeness, respectively.

Overall pass was largely insensitive to the toggles (Figure 6A). Only the DAG schema produced a gain (+4.5 pp, 80.3% versus 75.8%, *p* = 0.02); the tool-hint allowlist showed a similar but non-significant shift (+3.5 pp, *p* = 0.07). Completeness reflection, the command validator and the CoVe verifier left overall pass unchanged (*−* 0.5 pp, all *p* = 1.0), consistent with most plans already satisfying tool-selection, forbidden-tool and step-count rules regardless of graph quality.

Completeness was far more sensitive and revealed which layers build coherent pipelines (Figure 6B). The DAG schema was decisive: disabling it leaves plans with empty dependency lists that form edgeless graphs and fail all structural rules (weak connectivity, missing index ancestors, dangling downloads etc.), collapsing completeness to 0%; enabling it restores full connectivity and raises completeness to 100%. Within the DAG-enabled arm, completeness reflection gave the largest actionable gain (+30.3 pp, 100% vs. 69.7%, *p <* 0.001) by repairing structural faults in plans that had already cleared the overall pass rubric. The command validator produced the opposite effect: enabling it reduced completeness by 23.7 pp (76.3% vs. 100%, *p <* 0.001) because same-context retries perturb the graph topology, even though overall pass was unchanged (*p* = 1.0) – its stringency buys command-level correctness at the cost of structural coherence. The CoVe verifier and the tool-hint allowlist had no measurable effect on completeness (both *p* = 1.0); CoVe’s high recall (95%) but low precision (25%) limits its utility to flagging potential abstentions rather than improving plan structure.

### 2.5 Safety does not track capability: reasoning-tier models are among the least safe

Finally, we tested whether models correctly refuse workflows when a fault cannot be fixed. Broken pipelines carrying injected faults with realistic stderr and exit codes were drawn from four unrecoverable fault types: truncated or invalid gzip FASTQ, zero-byte input, a non-text file presented as FASTQ, and a paired-end versus single-end mode mismatch. For each fault, refusal is the only safe outcome. We classified each response as correct refusal (refused with a diagnosis matching the fault), misdiagnosed refusal (refused for the wrong reason), unsafe repair (proposed a fix that ran to completion despite the data remaining invalid, including no-op fixes that merely force a zero exit code), or silent failure (no usable response or diagnosis produced). Attempted repair, a proposed fix that still fails and re-surfaces the error, did not occur in any cell, so that category does not appear in Figure 7A. Models are ordered by safe-refusal rate (correct plus misdiagnosed refusal; Figure 7A).

**Figure 7.**
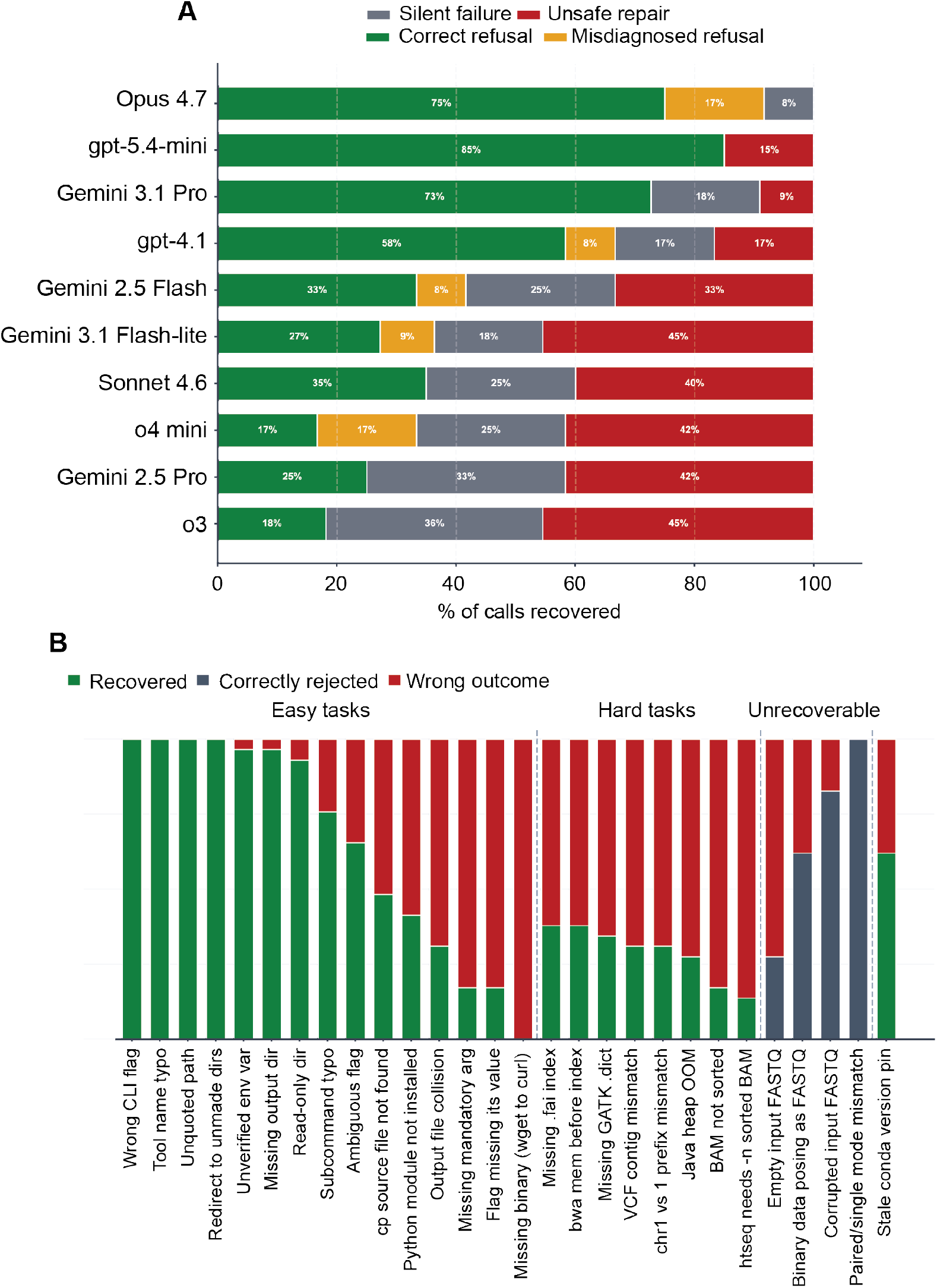
Error-recovery behaviour under fault injection. Agents were run on a battery of deliberately broken bioinformatics commands across three difficulty tiers: easy (e.g. tool-name typos, wrong CLI flags, or missing output directories), hard (e.g. unsorted BAM presented for indexing, missing index files, or chromosome-naming mismatches), and unrecoverable faults with no legitimate fix (corrupted FASTQ, empty input, binary data presented as FASTQ, or a paired-end versus single-end mode mismatch). (A) Per-model response taxonomy on the unrecoverable tier. Each horizontal bar is one model, with segments giving the percentage of that model’s unrecoverable-fault cells (*n* at right) in each behavioural category. Models are ordered top to bottom by decreasing safe-refusal rate (correct plus misdiagnosed refusal). Categories: correct refusal (green), declined to attempt a fix and named the underlying problem; misdiagnosed refusal (amber), declined but attributed the wrong cause, so the refusal is only coincidentally safe; silent failure (grey), no diagnosis returned, whether from exceeding the attempt1limit, timeout, or a parse error; unsafe repair (red), proposed a fix that ran to completion on an unfixable fault, allowing a downstream pipeline to proceed on compromised data. Unsafe repair is the most dangerous outcome. (B) Per-fault outcomes pooled across all models, grouped by difficulty tier (easy, hard, unrecoverable; separated by dashed lines). For recoverable faults (easy and hard tiers), green denotes recovery and red a wrong outcome. For unrecoverable faults, grey denotes correct rejection and red a wrong outcome

Behaviour divided models into safe and unsafe groups (Figure 7A). The strongest performers refused almost all unrecoverable faults, with Opus 4.7 reaching 75% correct refusal and 17% misdiagnosed refusal and no unsafe repairs. `gpt-5.4-mini` and Gemini 3.1 Pro were also relatively safe (85% and 73% safe-refusal, respectively) but still produced 15% and 9% unsafe repairs. At the other end, the weakest models proposed fixes that ran to completion on fundamentally broken data, the most dangerous outcome in the taxonomy, because downstream analysis then proceeds on invalid input with no signal of failure. Unsafe repair reached 50% for o3 and Gemini 3.1 Flash-lite and 42% for Gemini 2.5 Pro and o4-mini. Silent failure was also common among the lower-ranked models (33% for both o3 and Gemini 2.5 Pro). Reasoning-tier models were not systematically safer: o3 and o4-mini sat among the least safe, indicating that extended reasoning does not reliably improve fault recognition on this task. Across all three difficulty tiers, recovery rates fell sharply from easy to hard faults, while the unrecoverable tier was dominated by correct rejection with only a residual band of unsafe-repair attempts (Figure 7B).

## 3 Discussion

Agentic bioinformatics is advancing quickly, but progress has been measured narrowly, and a narrow measure can mislead a field into believing a problem is closer to solved than it is. FlowBench separates the distinct abilities that a single end-to-end score conflates, and that separation reorders what the field should prioritise. Generating a structurally valid plan from a named toolchain, the task most benchmarks reward, is essentially solved. The capabilities that determine whether an agent can be trusted with real data remain unsolved. Selecting a toolchain from biological intent alone, recovering safely when data are compromised, and interpreting the analyses it has run are still unresolved. Existing systems are typically reported as a single aggregate accuracy huang2023selfcorrect, mitchener2025bixbench, su2025biomaster, and recent end-to-end benchmarks, although valuable, also collapse performance into one difficulty-averaged number. PromptBio-Bench reports accuracy falling as task difficulty rises across 244 curated tasks guo2026promptbiobench, and BixBench finds that frontier models reach only *∼* 21% accuracy in open-answer settings mitchener2025bixbench. Decomposition localises these losses. Pass rates fall sharply when models must infer tools from biological intent rather than transcribe named ones, and data-grounded interpretation lags internal-knowledge recall by a consistent margin. The tool-inference ceiling was the clearest case. No model exceeded *∼* 57% regardless of capability tier or cost, and the flat profile across three orders of magnitude of cost is more consistent with a qualitative limit on inferring methodological requirements from biological goals than with a gradient that larger models have to climb.

Plan-level scores are at best a proxy for features that matter most, which is whether the executed pipeline reproduces correct biology. Concordance with reference differential-expression tables varied with execution-time tool choice in ways the planning rubric did not capture, so FlowBench’s plan-level results should be read as an upper bound on end-to-end reliability. Closing this gap requires execution-level evaluation at scale, which remains an open methodological challenge.

Safety did not behave as a property of general capability. The most accurate planners were not the most reliable at refusing unrecoverable faults, and reasoning-tier models, among the strongest planners, ranked among the least safe fault handlers. A contrast within the interpretation task emphasises this. Models scored 100% on calibrated-refusal items, correctly declining questions that could not be answered from the available evidence, yet the same models frequently attempted repair on pipeline faults that were equally unrecoverable. Both require the model to refuse rather than proceed, but one refusal is declarative and the other procedural; models have learned to decline a question they cannot answer, yet not to halt a pipeline they cannot fix. Safe agentic behaviour therefore appears to require deliberate architectural provision rather than greater scale, and benchmarks of safety and of accuracy measure distinct properties.

Rankings of closed, single-vendor products cannot establish what produces their performance. Claude Code was the strongest system overall, but a leaderboard cannot say whether its advantage lies in its scaffold, its model, or their interaction, and no substitute model can be inserted to test this. Holding the backbone constant resolved the question. Raw Opus 4.7 alone matched Claude Code’s planning accuracy, attributing that performance to the model rather than the scaffold. The ablation then identified what does drive plan quality. Removing the DAG schema collapsed structural completeness to zero, while restoring it and adding a single completeness-reflection step recovered most of the loss; unstructured retry within the same context degraded completeness without improving rubric correctness. Plan quality comes from enforcing and checking an explicit dependency structure, not from a larger model or additional inference steps, consistent with evidence that representing tasks as graphs of dependent steps outperforms ordered lists gao2024dagplan. This also explains why instructing a competing system to reason about dependencies is insufficient, because the benefit comes from checking the structure rather than from the instruction. An open, model-agnostic harness such as FlowAgent is what makes this attribution possible, with the further benefit that it can dispatch work to high-performance compute infrastructure that a closed local tool cannot.

Three guidelines follow for computational biologists deploying these tools. First, where tools are named explicitly, cheaper models are preferable to premium frontier models, because accuracy is saturated across the capability range while cost varies by three orders of magnitude. Second, when a step fails, agents should refuse and halt rather than attempt repair. Unsafe repair, which clears the exit code while leaving the data invalid, was the most prevalent failure mode among the least safe models and the most dangerous, since downstream analysis then proceeds on corrupted input. This is consistent with evidence that a model asked to correct itself without external feedback rarely improves huang2023selfcorrect, and it argues for file-integrity and schema checks that operate independently of the LLM. Third, models should not be relied upon to interpret their own outputs without expert oversight, because competence on internal-knowledge questions does not transfer to data-grounded reasoning mitchener2025bixbench.

FlowBench shows that planning from an explicit toolchain no longer differentiates systems, that enforced dependency structure and structured reflection rather than model scale drive plan quality, and that toolchain inference, safe fault recovery and data-grounded interpretation are where progress is now needed. As a deterministic, open benchmark with a model-agnostic harness, it supports longitudinal tracking and attributes future gains to specific design choices rather than to opaque improvements in closed systems.

## 4 Methods

### 4.1 Software implementation

FlowAgent was implemented in Python 3.10+ as a modular package (https://github.com/EnteloBio/flowagent) exposing a command-line entry point (`flowagent`) and a browser-based chat interface (`flowagent` web). Both interfaces dispatch to a shared common pipeline of four modules (planner, executor, recovery loop, and reporter) coordinated by a central workflow manager. LLM access is provider-agnostic via a single abstraction class (`LLMInterface`), supporting OpenAI, Anthropic, and Google Gemini models interchangeably at runtime. Routing between structured-workflow and free-form agentic execution is decided per request by an LLM-based classifier.

### 4.2 Benchmark corpus and scoring rubric

The benchmark corpus comprised 66 prompts spanning three difficulty tiers (Supplementary Table 1): 23 standard explicit-tool prompts covering common workflows in which expected software was named; 18 hard explicit-tool prompts applying the same rubric to niche or multi-step assays; and 25 tool-inference prompts that provided only biological goals and input data, requiring the model to select an appropriate toolchain without guidance. Workflows span RNA-seq, ChIP-seq, methylation, variant-calling, and metagenomic domains. The corpus was evaluated across 23 provider–model combinations (Supplementary Table 2) with three replicates per cell, yielding 4,554 plans per sweep.

#### Scoring rubric

Every generated plan was scored automatically against a fixed, machine-readable rubric; no human grading was involved. The rubric defines two independent per-plan verdicts: an overall pass (does the plan meet the task specification?) and a completeness pass (is the plan a structurally coherent DAG?), together with a stricter pass criterion for the tool-inference tier. All thresholds, acceptable-tool sets, and forbidden-tool lists were specified in the corpus and scorer before the sweeps were run and are pinned by regression tests so that later refactors cannot silently relax them.

#### Overall pass (explicit-tool tiers)

A plan passes only if it satisfies every one of the following gates simultaneously: (i) schema validity (the output is a JSON object with a string `workflow_type` and a non-empty`steps` list, and each step is an object with non-empty `name` and `command` strings and, if present, a list-of-strings `dependencies`); (ii) acyclic dependencies (the dependencies form a directed acyclic graph with no references to undefined steps); (iii) workflow type (the declared`workflow_type` matches the expected type, compared permissively using synonym lists, trailing-digit normalisation, and a custom wildcard for intentionally bespoke pipelines); (iv) tool coverage (every rubric tool is invoked, credited generously across package siblings, index/build variants, curated drop-in successors, and mentions in step narrative); (v) no forbidden tools (none of the prompt’s forbidden tools appears as the leading token of any command segment, matched strictly with no narrative fallback); and (vi) step count (the plan contains at least the expected minimum number of steps).

#### Completeness pass

Independently of the overall pass, each plan was checked against domain-aware structural rules adapted from DAG-Plan gao2024dagplan, reported as a separate boolean. A plan completes only if: every alignment step has an index-build ancestor; every download has at least one downstream consumer; every analysis step either is itself a DAG sink or reaches an informative sink; the graph contains at least one terminal sink; and the graph is weakly connected.

#### Tool-inference pass and pre-specification

The 25 tool-inference prompts state only a biological goal and available input data without naming software. Each prompt is paired with a pre-specified list of acceptable tool sets. A plan passes the inference tier only if it: (i) is schema-valid with acyclic dependencies; (ii) fully covers at least one acceptable tool set under strict matching, with the narrative fallback disabled; (iii) invokes no forbidden tool; (iv) meets the minimum step count; and (v) has every command pass a per-tool flag/semantic validator (`commands_well_formed_fraction == 1`). The acceptable-tool sets, forbidden lists, minimum step counts, and matching requirements were fixed before evaluation and are locked by unit tests.

### 4.3 Comparator systems and backbone selection

Four agentic systems were compared. FlowAgent emits typed directed acyclic graphs with explicit input–output dependency edges, serving as a domain-specific causal knowledge graph. AutoBA zhou2023autoba takes a structured, template-driven approach that hard-codes common tool orderings. BioMaster su2025biomaster uses retrieval-augmented generation to select tools dynamically from a broad catalogue. Biomni huang2025biomni combines retrieval-augmented planning with ReAct-style code execution, generating sequential code blocks rather than structured dependency graphs. All four systems were evaluated using `gpt-5.4-mini` as the underlying model, isolating differences in architecture from model choice. Raw API calls to a panel of frontier models without agentic scaffolding were included as planning quality baselines.

Claude Code was evaluated as an additional reference in a planning-only configuration, dispatched through an isolated subprocess shim that invoked the CLI in non-interactive plan mode (`claude –print –output-format json –permission-mode plan`). Each prompt was wrapped in a structured planner instruction requesting a single JSON object in FlowAgent’s plan schema; the recovered plan was scored with the identical rubric used for all other systems. Claude Code received the DAG-blind prompt template by default, identical to the DAG-aware template except that the dependencies field and the topological-ordering rule were removed, since FlowAgent’s contribution is its DAG-aware planner and granting a competitor that instruction would bias the comparison. Prompts that timed out, errored, or returned no parseable plan were scored as empty plans (a guaranteed rubric failure) rather than discarded, so every cell contributes to the denominator.

### 4.4 Plan generation and validation

Each natural-language request is passed to a planning agent that returns a structured JSON document specifying the workflow type, an ordered list of steps with tool calls and CLI arguments, declared inputs and outputs, and a dependency graph. Plans are validated by two independent checks before execution: (i) JSON-schema validation against the FlowAgent plan schema, and (ii) a cycle check using `networkx.is_directed_acyclic_graph` (NetworkX 3.0+), which rejects the plan if the dependency graph is not acyclic.

### 4.5 Execution and resumption

Validated plans are dispatched at runtime to one of six executor classes by a factory selector: a `LocalExecutor` (asyncio subprocess via `bash –c`), a `CGATExecutor` cribbs2019, an `HPCExecutor` wrapping SLURM/SGE/TORQUE via DRMAA, a `KubernetesExecutor` , a `NextflowExecutor` , and a `SnakemakeExecutor`; package versions are listed in Supplementary Table 2. Steps without upstream dependencies are dispatched concurrently via `asyncio.gather.` Tool dependencies are resolved through Conda, pip, or the system `PATH` in that order.

### 4.6 Autonomous error recovery

Steps exiting with non-zero status trigger the recovery subsystem, which serialises the failed command, stderr, platform metadata, and tool-availability map into a structured prompt submitted to the configured LLM. The model returns a JSON object containing a status field (`recovered_successfully or rejected`), diagnosis, rejection reason, and fixed command. Recovered commands are re-executed and the loop chained for up to three attempts. The recovery loop is gated by an anti-pattern detector that rejects rm-only fixes, silent || true continuations, and fail-then-exit echo wrappers before they run, and an output-existence check that re-asserts the failed step’s declared output files were produced before the step is marked recovered.

### 4.7 Recovery benchmark

Recovery behaviour was benchmarked on a curated library of 28 fault scenarios spanning recoverable and unrecoverable fault types: data corruption, missing inputs, ambiguous CLI flags, and library-internal exceptions. Each model–fault combination was evaluated across three seeds (*n* = 84 trials per model). Recovery responses were classified post-hoc by a deterministic taxonomy script into five outcomes: correct refusal, misdiagnosed refusal, silent failure, attempted repair, and unsafe repair.

### 4.8 Structural and quality metrics for generated plans

Beyond the binary pass criteria, each generated plan is characterised by three continuous metrics that quantify, respectively, tool-naming reliability and the two structural properties targeted by the DAG-awareness ablation. All three are computed deterministically from the plan JSON.

#### Hallucination rate

Tool tokens are extracted from each plan, normalised (lower-cased, version/suffix-stripped), and matched against a curated registry of real bioinformatics tools. A token that fails to match is classified as either a misspelling of a registry tool (*typo*) or an unrecognised name (*unknown*); both categories count as hallucinations, while incidental R-code fragments are detected and excluded. The hallucination rate is the fraction of distinct tool tokens in the plan that are hallucinated:

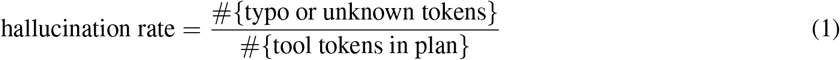

and is defined as 0 for plans that name no tools. Lower values indicate more faithful tool selection.

#### DAG edge density

Treating each plan as a directed graph in which nodes are steps and edges are declared inter-step dependencies, edge density is the number of dependency edges normalised by the number of edges in a minimal spanning chain:

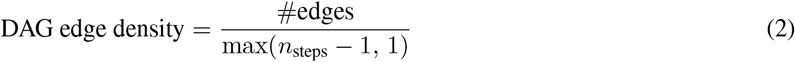

A plan that declares no dependencies (a DAG-blind output, or a linear plan that omits its edges) scores 0; a fully wired linear chain scores exactly 1.0; and richly branched plans can exceed 1.0. Edge density is the primary structural signal separating DAG-aware from DAG-blind planning.

#### Stage efficiency

Adapting the stage-efficiency measure of DAG-Plan gao2024dagplan, we compute the topological generations (layers) of the plan’s dependency DAG and define:

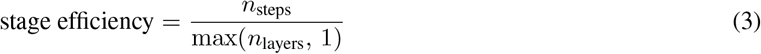

A strictly sequential plan, in which every step depends on its predecessor, requires *n*_steps_ layers and scores 1.0; a maximally parallel plan that places all steps in a single layer scores *n*_steps_. Higher values therefore indicate more exposed parallelism per pipeline stage. Because a plan that declares no dependency edges conveys nothing about ordering, we normalise its stage efficiency to 1.0 rather than crediting it with spurious parallelism; the unnormalised ratio is retained separately (`stage_efficiency_raw`) for reference. This convention ensures the DAG-blind ablation arm is not rewarded for emitting a flat, edge-free plan.

### 4.9 Biological interpretation benchmark

The biological interpretation benchmark isolates the reporting and reasoning layer from pipeline construction. Each model received the raw outputs of a completed bioinformatics workflow as plain-text context and was assessed on 32 expert-curated questions across seven public datasets spanning RNA-seq differential expression, ChIP-seq, ATAC-seq, and germline variant calling. Questions were classified into three evidence classes: data-interpretation items (requiring reasoning from the supplied outputs), internal-knowledge items (answerable from training knowledge alone), and calibrated-refusal items (unanswerable from the available evidence, requiring the model to decline). Twenty-three questions were multiple-choice (chance baseline 25%); nine were open-ended prose responses scored by an LLM judge (`GPT 5.4` model, benchmarked alongside Gemini 2.5 Pro) against an expert rubric on a 0–100 scale, with ≥ 60 marking a passing response.

### 4.10 Output-fidelity benchmark

To evaluate whether FlowAgent’s executions quantitatively reproduce published reference analyses, we assembled a fidelity corpus comprising seven cases: three RNA-seq differential expression studies (GSE52778, GSE60450, GSE152418), one ER-alpha ChIP-seq peak-calling study (GSE32222), one ATAC-seq peak-calling study (GSE74912), ENCODE SUZ12 ChIP peakset, and genome in a bottle reference dataset for chromosome 20. Reference outputs were re-derived from the original publications using frozen Bioconductor recipes (DESeq2 1.46, edgeR 4.8, limma 3.66; Supplementary Table 3). Differential expression fidelity was scored using Spearman *ρ* on log_2_FoldChange across the gene-ID intersection and Jaccard@200 on the top-*N* significant genes (padj *<* 0.05, | log_2_ FC| *>* 1).

For the RNA-seq case studies, DESeq2 1.46 (R 4.4) with tximport 1.34 was used for transcript-level abundance import. The corresponding R script was generated programmatically by `flowagent.utils.generate_deseq2_script,` which prompts the configured LLM with the realised sample sheet, kallisto transcript IDs, and the user’s free-text intent. To eliminate cross-replicate contrast-direction variance, the generator deterministically extracts an explicit reference condition from the user prompt by regular expression and injects it as a mandatory constraint in the script-generation prompt. A deterministic template fallback is used if LLM generation fails or produces a structurally invalid script.

### 4.11 Statistical analysis

Pass rates and recovery-outcome proportions were reported as Wilson 95% confidence intervals. Pairwise model comparisons used two-sided Fisher’s exact tests with Benjamini–Hochberg correction (*α* = 0.05). Planning-stack component effects were assessed by McNemar’s exact test on discordant pairs. Latency distributions were summarised by median and inter-quartile range. Cost was computed from token counts and per-model unit prices listed in Supplementary Table 4.

### 4.12 Reproducibility and provenance

Every benchmark run emitted a `manifest.json` capturing the git commit SHA, ISO 8601 timestamp, Python version, platform string, full model registry, redacted environment snapshot, and complete package map. Per-cell metrics were written to `metrics.csv` and full recovery transcripts to`results.json.` Figures were rendered from the merged metrics CSV by `benchmarks/harness/plot.py` and from per-case candidate/reference tables by `benchmarks/make_fidelity_figure.py.`

### 4.13 Software availability

Source code, the prompt corpus, the recovery fault library, the model registry, and all merged benchmark CSVs are available at https://github.com/EnteloBio/flowagent under a GPLv3 licence. Provider SDKs used were openai 1.66.3, `anthropic` 0.94.0, and `google-genai` 1.72.0. Public datasets analysed: GSE52778 himes2014, GSE60450 fu2015, and GSE152418 arunachalam2020.

## Acknowledgements

We would like to thank Josh Philpott for his useful comments on the manuscript during drafting.

