## Supplementary Figures for "FlowBench: separating planning, fault recovery and interpretation in agentic bioinformatics"

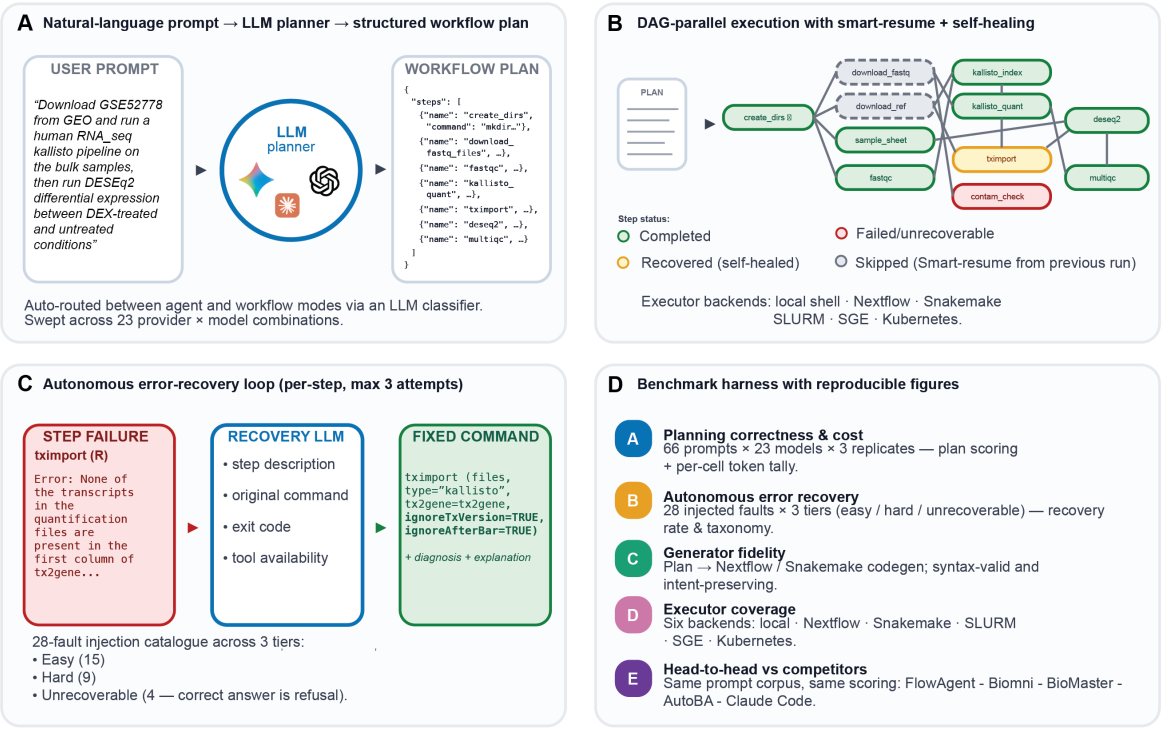


**Supplementary Figure 1. FlowAgent architecture and the FlowBench harness.** (A) The planner maps a natural-language request to a structured JSON workflow plan, with routing between agentic and structured-workflow execution decided per request by an LLM classifier; plans were swept across 23 provider and model combinations. (B) Validated plans are executed as a directed acyclic graph, with independent steps dispatched in parallel, per-step checkpointing for smart resume, and an LLM recovery loop. Step status is shown as completed, recovered (self-healed), failed or unrecoverable, and skipped on resume, across six executor backends (local shell, Nextflow, Snakemake, SLURM, SGE and Kubernetes). (C) The autonomous recovery loop runs for up to three attempts per failed step; the recovery model receives the step description, original command, exit code and tool-availability map, and returns a fixed command with a diagnosis. The injected-fault catalogue comprises 28 scenarios across three tiers (15 easy, 9 hard and 4 unrecoverable, for which refusal is the correct outcome). (D) The benchmark harness scores planning correctness and cost, autonomous error recovery, generator fidelity, executor coverage, and head-to-head comparison against competitor systems, and renders the figures reproducibly.

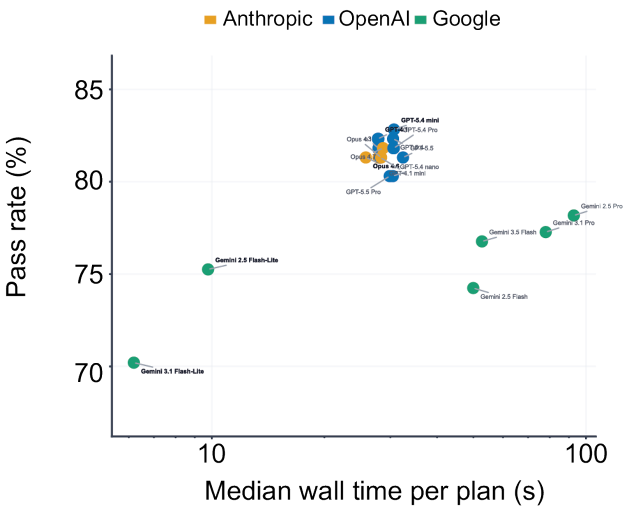


**Supplementary Figure 2. Planning accuracy against latency across 23 models.** Overall pass rate (%) against median wall-clock time per plan (s, log scale), one point per model, coloured by provider (Anthropic, OpenAI and Google). Anthropic and OpenAI models cluster at high pass rate (approximately 80 to 83%) and low to moderate latency, whereas Google models span a wider latency range at lower pass rates; the two fastest models (Gemini 3.1 and 2.5 Flash-Lite) are also the least accurate.

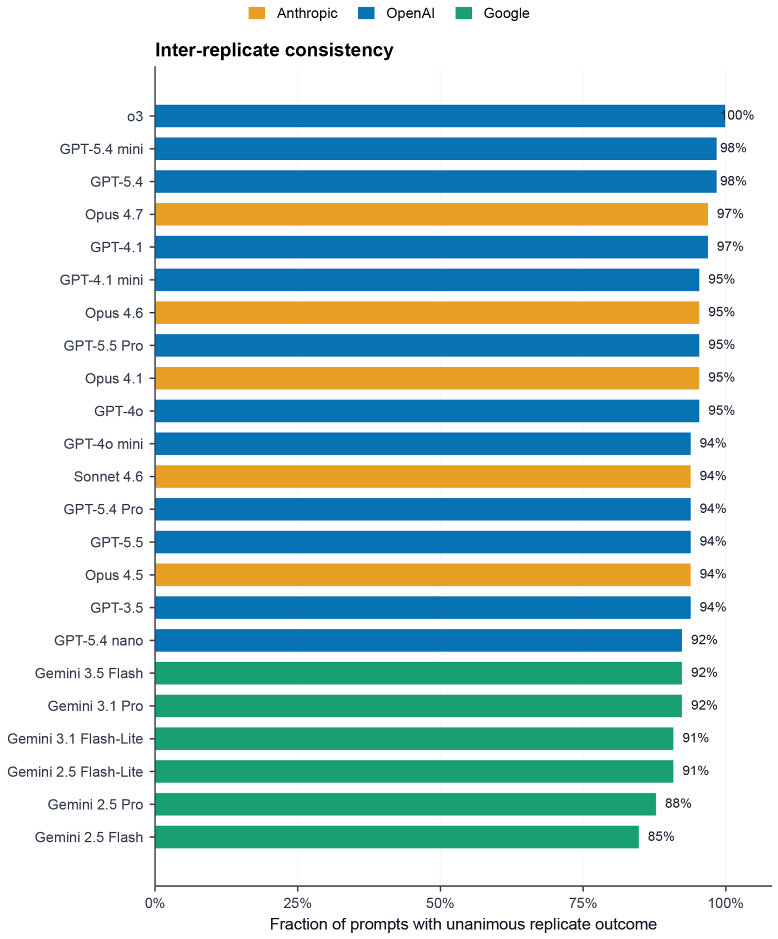


**Supplementary Figure 3. Inter-replicate consistency of planning outcomes.** For each model, the fraction of the 66 prompts whose three replicates returned an identical pass or fail outcome, ordered by decreasing consistency and coloured by provider. Values range from 100% (o3) to 85% (Gemini 2.5 Flash), indicating that triplicate scoring is stable and that the reported pass-rate differences are not driven by replicate noise.

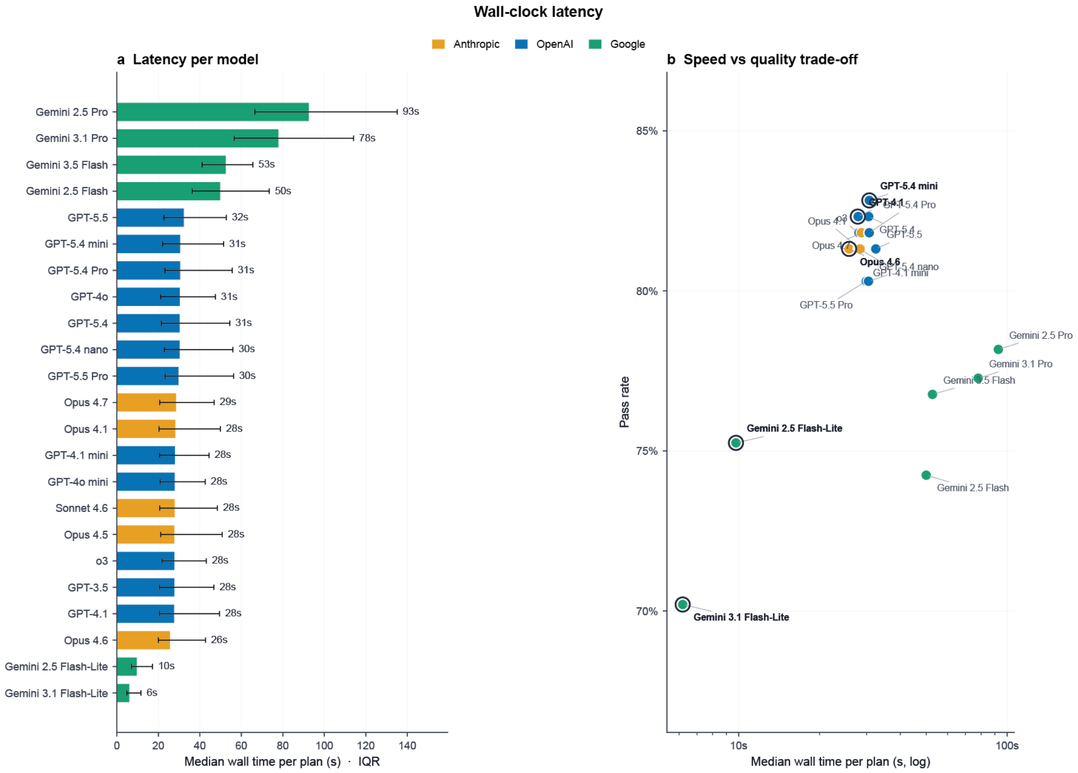


**Supplementary Figure 4. Wall-clock latency across 23 models.** (a) Median wall-clock time per plan (s) with interquartile range, ordered slowest to fastest and coloured by provider; Google Pro models are slowest (Gemini 2.5 Pro, 93 s) and the two Flash-Lite models fastest (6 to 10 s). (b) Pass rate against median wall time (log scale), with the fastest model per provider ringed. Latency varies by roughly an order of magnitude with no corresponding gain in pass rate.

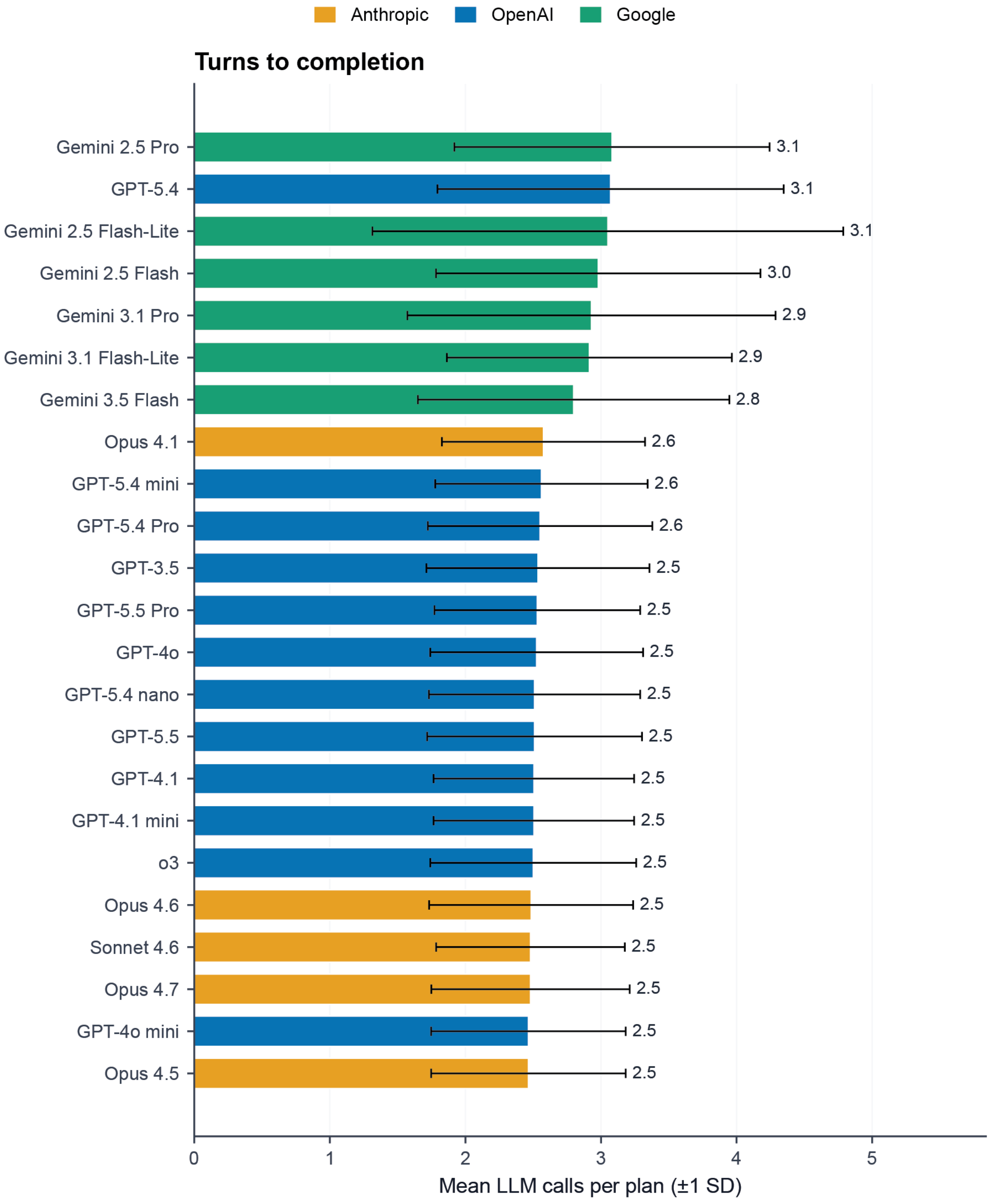


**Supplementary Figure 5. Planner round-trips per plan.** Mean number of LLM calls required to produce a plan (±1 SD) for each model, ordered descending and coloured by provider. Google models average the most calls (2.8 to 3.1) and Anthropic and OpenAI models slightly fewer (approximately 2.5; with one gpt-5.4 outlier), a modest, provider-linked difference in planning verbosity.

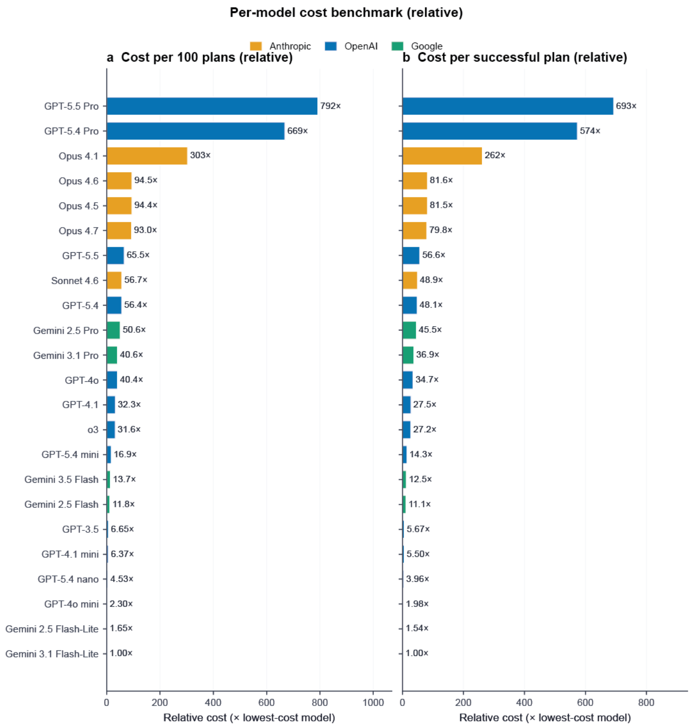


**Supplementary Figure 6. Relative planning cost across 23 models.** (a) Cost per 100 plans and (b) cost per successful plan, each normalised to the cheapest model (Gemini 3.1 Flash-Lite, 1.00×) and coloured by provider. Cost spans roughly three orders of magnitude, from 1.00× to 792× per 100 plans and to 693× per successful plan, with the reasoning-tier OpenAI models (GPT-5.5 Pro, GPT-5.4 Pro) and Opus 4.1 the most expensive.

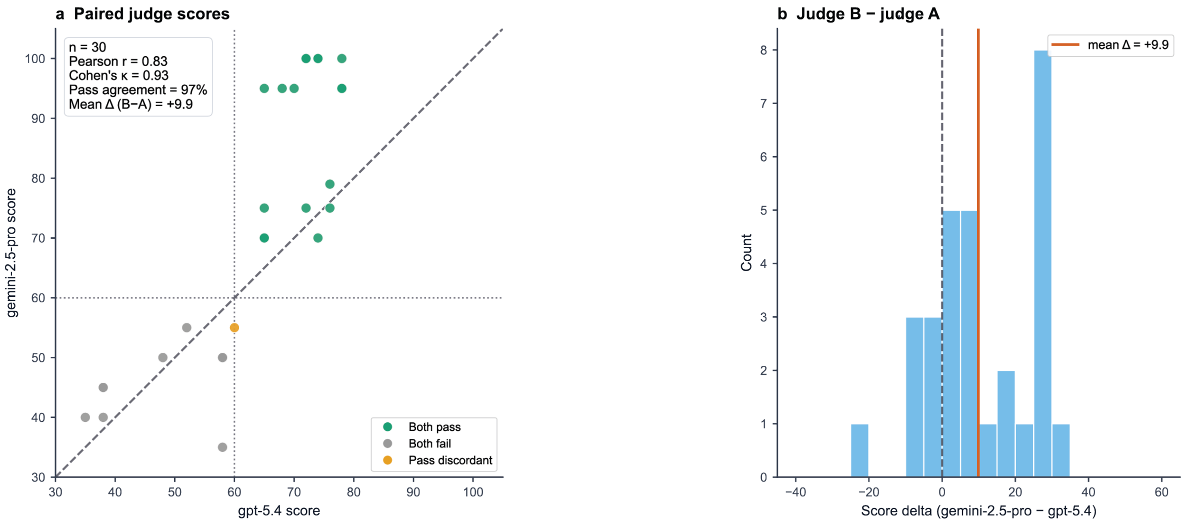


**Supplementary Figure 7. Inter-judge calibration on open-ended interpretation responses.** Agreement between the primary judge (gpt-5.4, judge A) and a second judge (gemini-2.5-pro, judge B) over n = 30 open-ended answers. (a) Paired judge scores (0 to 100); the dashed line marks y = x and the dotted lines the passing threshold of 60, with points classified as both judges passing, both failing, or pass-discordant. The judges agree closely (Pearson r = 0.83, Cohen's κ = 0.93, 97% pass agreement). (b) Distribution of the per-answer score difference (judge B minus judge A); judge B scores 9.9 points higher on average, a near-uniform offset that does not change the pass or fail classification.

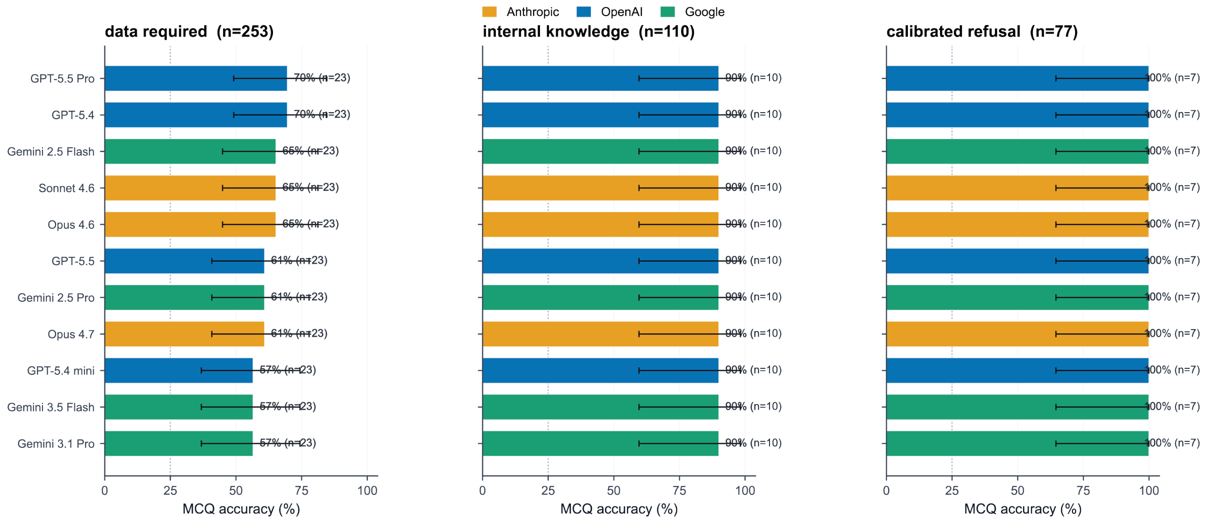


**Supplementary Figure 8. MCQ accuracy by evidence class.** Multiple-choice accuracy (%) for 11 models, stratified by the evidence each question requires (chance = 25%), coloured by provider and shown with Wilson 95% confidence intervals: data required (n = 253), internal knowledge (n = 110) and calibrated refusal (n = 77). Accuracy is uniformly high on internal-knowledge questions (90%) and on calibrated-refusal items (100%), but falls to 57 to 70% when the answer must be derived from the supplied data, isolating data-grounded reasoning as the limiting evidence class.

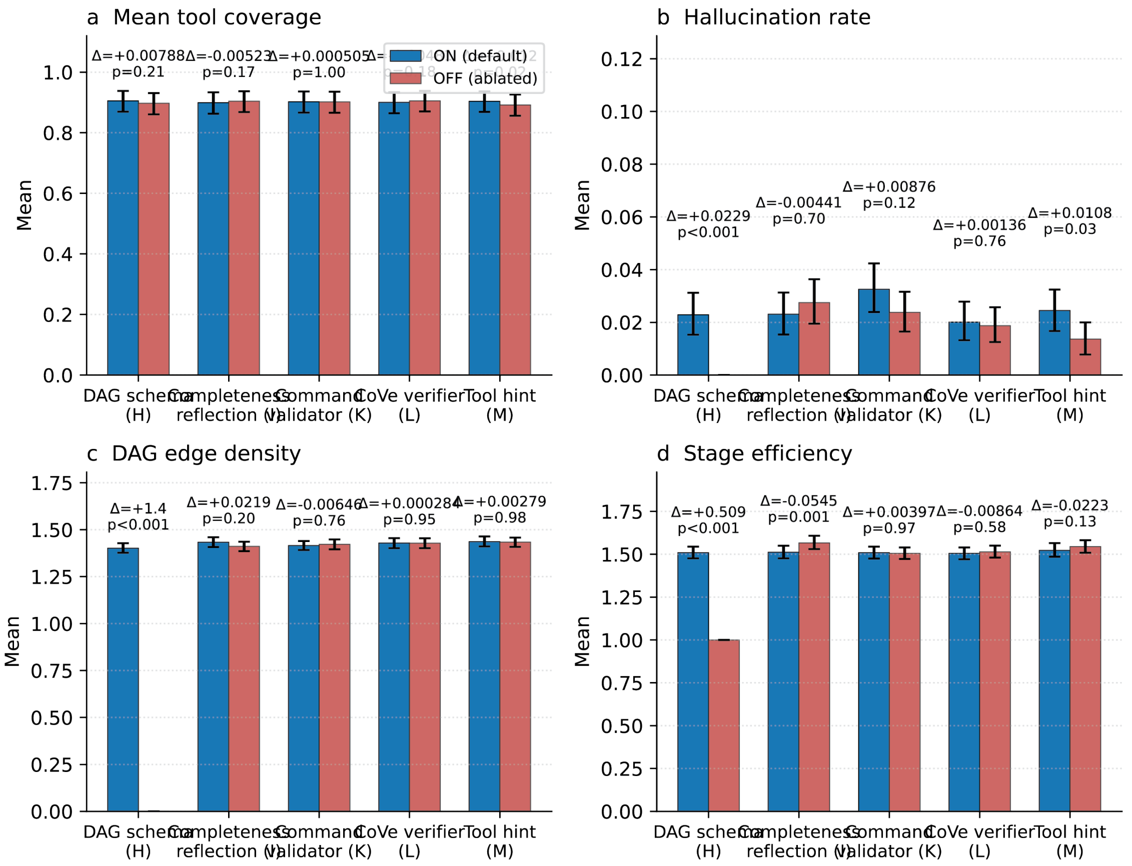


**Supplementary Figure 9. Per-component paired ON/OFF ablation of the FlowAgent planning stack.** Each of the five planning-stack components (DAG schema, completeness reflection, command validator, CoVe verifier and tool-hint allowlist) was toggled on (default, blue) and off (ablated, red) while the others were held at default (gpt-5.4-mini, 66 prompts × 3 replicates), with each ON/OFF pair annotated by Δ (ON minus OFF) and its paired p-value. Panels report (a) mean tool coverage, (b) hallucination rate, (c) DAG edge density and (d) stage efficiency. Tool coverage is near-ceiling and insensitive to every toggle. The DAG schema is the dominant structural driver; ablating it collapses edge density to zero (Δ = +1.4, $p < 0.001$) and reduces stage efficiency from approximately 1.5 to 1.0 (Δ = +0.509, $p < 0.001$). The command validator is the only component whose removal raises stage efficiency (Δ = −0.0545, $p = 0.001$), consistent with same-context retries perturbing graph topology.
